# HDA6 regulates chloroplast biogenesis and photosynthesis in Arabidopsis through deacetylation of non-histone proteins

**DOI:** 10.1101/2025.10.28.685251

**Authors:** Hao Wang

**Author notes:** **Corresponding Author:** Hao Wang, Ph.D., Professor.

## Abstract

Histone deacetylase 6 (HDA6) plays a multifaceted role in plant development, yet its function in chloroplast biogenesis remains largely unexplored. Here, we show that the *axe1-4* loss-of-function mutant exhibits pale-green leaves, reduced chlorophyll content, and impaired photosynthetic efficiency, accompanied by smaller and morphologically distorted chloroplasts. Transcriptomic analyses revealed global down-regulation of chloroplast-associated genes, including those involved in chlorophyll biosynthesis and thylakoid organization. Blue-native PAGE analyses of *axe1-4* leaves demonstrated markedly decreased accumulation of PSI, PSII, and LHCII complexes, while cryo-electron tomography confirmed disorganized thylakoid architecture characterized by swollen membranes and disrupted grana stacking. Confocal imaging of *pUBQ::HDA6-GFP* plant leaves indicated both nuclear and chloroplast localization, suggesting a dual role in chromatin regulation and organellar protein deacetylation. Together, these findings identify HDA6 as a pivotal epigenetic regulator integrating transcriptional control with photosynthetic machinery assembly and chloroplast development in *Arabidopsis thaliana*.

## Introduction

Chloroplasts are the photosynthetic organelles of plant cells and play indispensable roles in energy conversion, carbon fixation, and numerous biosynthetic processes essential for plant growth and productivity. Their biogenesis is a highly coordinated process that requires intricate communication between nuclear and plastid genomes, ensuring proper expression and assembly of photosynthetic complexes, pigments, and membrane structures (Daum and Kühlbrandt, 2011; Pribil et al., 2014; Rast et al., 2015; Komenda et al., 2024). Perturbations in this coordination often result in defective chloroplast differentiation and impaired photosynthetic performance, as exemplified by mutants disrupted in nucleo-cytoplasmic signaling, protein import, or thylakoid organization (Jarvis and López-Juez, 2013; Pribil et al., 2014). While transcriptional and translational regulation of chloroplast development has been extensively studied, emerging evidence suggests that epigenetic mechanisms also play pivotal roles in modulating plastid biogenesis through nuclear gene expression and post-translational control.

Histone acetylation and deacetylation represent key epigenetic modifications that dynamically regulate chromatin structure and gene transcription (Wu *et al*., 2008; Chen *et al*., 2020; Zhang *et al*., 2024). In plants, histone deacetylases (HDACs) are classified into three major families including RPD3/HDA1, SIR2, and HD2, among which members of the RPD3-type HDACs are the best characterized (Chen *et al*., 2020; Zhang *et al*., 2024). These enzymes catalyze the removal of acetyl groups from lysine residues on histones and a variety of non-histone substrates, thereby influencing transcriptional activity, RNA processing and protein stability (Jin *et al*., 2024). In *Arabidopsis thaliana*, HDA6 has emerged as a multifunctional regulator that interacts with DNA methyltransferase MET1 and histone demethylases LDL1/2 and FLD to modulate flowering time, transposon silencing, circadian rhythm, and stress responses (Liu *et al*., 2012; Martignago *et al*., 2019). Beyond its nuclear role, recent reports have revealed that several HDACs, such as AtHDA14 and OsHDA710, localize to chloroplasts and mitochondria to deacetylate metabolic enzymes, suggesting broader, non-canonical roles in organellar function (Finkemeier *et al*., 2011). Nevertheless, the contribution of HDA6 to chloroplast development and photosynthetic efficiency has not been elucidated.

Chloroplast proteomic studies have demonstrated extensive lysine acetylation on photosynthetic enzymes and thylakoid proteins, including Rubisco activase, ATP synthase, and light-harvesting complex subunits, implying regulatory functions of reversible acetylation in photosystem assembly and energy conversion (Finkemeier *et al*., 2011). However, whether histone deacetylases directly participate in maintaining chloroplast protein homeostasis and membrane organization remains largely unknown. Given that HDA6 integrates multiple chromatin and transcriptional pathways in the nucleus, we hypothesized that it might also coordinate nuclear-encoded chloroplast gene expression with organellar proteostasis through histone and non-histone deacetylation.

In this study, we investigated the role of HDA6 in chloroplast development using the *axe1-4* loss-of-function mutant of *Arabidopsis thaliana*. The *axe1-4* plants exhibited a pale-green phenotype, reduced chlorophyll content, and aberrant chloroplast morphology, indicative of defective photosynthetic differentiation. Transcriptomic profiling revealed widespread down-regulation of chloroplast-associated genes involved in pigment biosynthesis and thylakoid organization. Blue-native gel and proteomic analyses showed reduced accumulation of PSI, PSII, and LHCII complexes, while cryo-electron tomography demonstrated severe disorganization of thylakoid membranes and grana stacking. Furthermore, confocal microscopy of *pUBQ::HDA6-GFP* plants revealed both nuclear and chloroplast localization of HDA6, supporting its dual roles in epigenetic regulation and non-histone deacetylation.

## Results and Discussion

### HDA6 is essential for chloroplast development and photosynthetic efficiency

To elucidate the physiological significance of HDA6 in chloroplast biogenesis, we analyzed the *axe1-4* loss-of-function mutant. Figure 1A depicts the schematic structure of the HDA6 protein, highlighting the catalytic histone deacetylase domain and the aspartate-rich C-terminal region in which the *axe1-4* mutation occurs. Morphologically, *axe1-4* plants exhibited a reduced rosette diameter, thinner leaves, and a pale-green coloration relative to the wile type (WT) (Figure 1B). Quantitative pigment analysis confirmed significant decreases in chlorophyll a and b contents (Figure 1C), indicating impaired photosynthetic capacity. Confocal imaging of chlorophyll autofluorescence revealed smaller and misshapen chloroplasts in *axe1-4* mesophyll cells (Figure 1D), and statistical assessment of projected area demonstrated a highly significant reduction compared with the wild type (Figure 1E).

**Figure 1.**
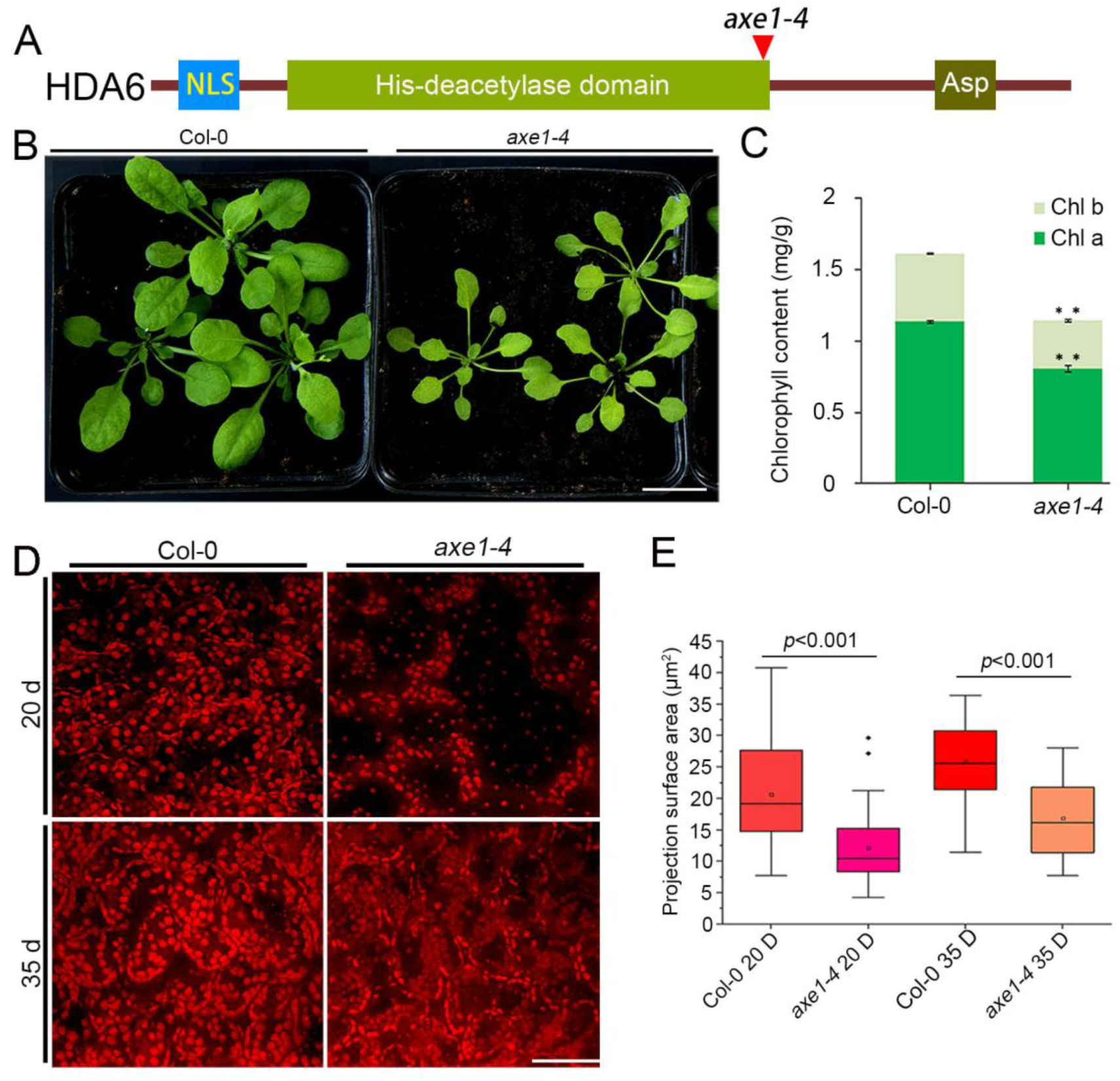
HDA6 is required for chloroplast development and photosynthetic efficiency in Arabidopsis. (A) Schematic representation of the HDA6 protein domain structure showing the nuclear localization signal (NLS), histone deacetylase (His-deacetylase) domain, and the C-terminal aspartate-rich (Asp) region. The position of the *axe1-4* mutation is indicated by a red arrowhead. (B) Growth phenotypes of 20-day-old wild-type (WT) and *axe1-4* mutant plants grown under identical conditions. The *axe1-4* mutant exhibits reduced leaf size and pale-green coloration. (C) Quantification of chlorophyll content in WT and *axe1-4* plants. Both chlorophyll a and b levels are significantly decreased in *axe1-4*. Data represent mean ± SD (n = 3). Asterisks indicate significant differences compared with WT (*p* < 0.01, Student’s *t*-test). (D) Chlorophyll autofluorescence images of mesophyll cells from 20-day-old and 35-day-old plants showing chloroplast morphology. The *axe1-4* mutant displays smaller and irregularly shaped chloroplasts compared with WT. Scale bar = 10 µm. (E) Quantification of chloroplast projection surface area based on confocal images shown in (D). Boxplots indicate median values, interquartile ranges, and outliers; circles represent individual data points. Statistical significance was determined using Student’s *t*-test (*p* < 0.001).

Such phenotypes parallel those observed in chloroplast biogenesis mutants such as *pp7l* and *sco3*, in which defective nucleo-cytoplasmic signaling leads to delayed chloroplast differentiation and abnormal thylakoid organization (Xu *et al*., 2019). The pale-green phenotype of *axe1-4* suggests that HDA6 is necessary not only for pigment accumulation but also for the developmental progression of proplastids into functional chloroplasts. Considering that HDA6 belongs to the RPD3-type HDAC family-enzymes that remove acetyl groups from lysine residues of histones and non-histone substrates. Its mutation likely disrupts chromatin accessibility and transcriptional programs required for plastid biogenesis. This interpretation is consistent with prior findings that histone acetylation balance, mediated by HDACs such as HDA6 and HDA19, plays a fundamental role in plant organ development and metabolic reprogramming (Wang *et al*., 2014). The reduced chlorophyll content and chloroplast deformities in axe1-4 therefore reflect a broader failure in photosynthetic differentiation, positioning HDA6 as a core regulator linking epigenetic control with chloroplast morphogenesis.

### Loss of HDA6 alters transcriptional regulation of chloroplast-associated genes

To explore the molecular basis of the chloroplast defects, transcriptome profiling was conducted. The volcano plot in Figure 2A shows extensive transcriptional reprogramming in *axe1-4*, with a large set of chloroplast-associated genes significantly down-regulated. Gene Ontology (GO) enrichment analysis highlighted over-representation of terms corresponding to chloroplast envelope, thylakoid membrane, and stroma components (Figure 2B). Notably, transcripts encoding enzymes involved in chlorophyll biosynthesis and structural factors critical for chloroplast organization were markedly suppressed (Figure 2C and D).

**Figure 2.**
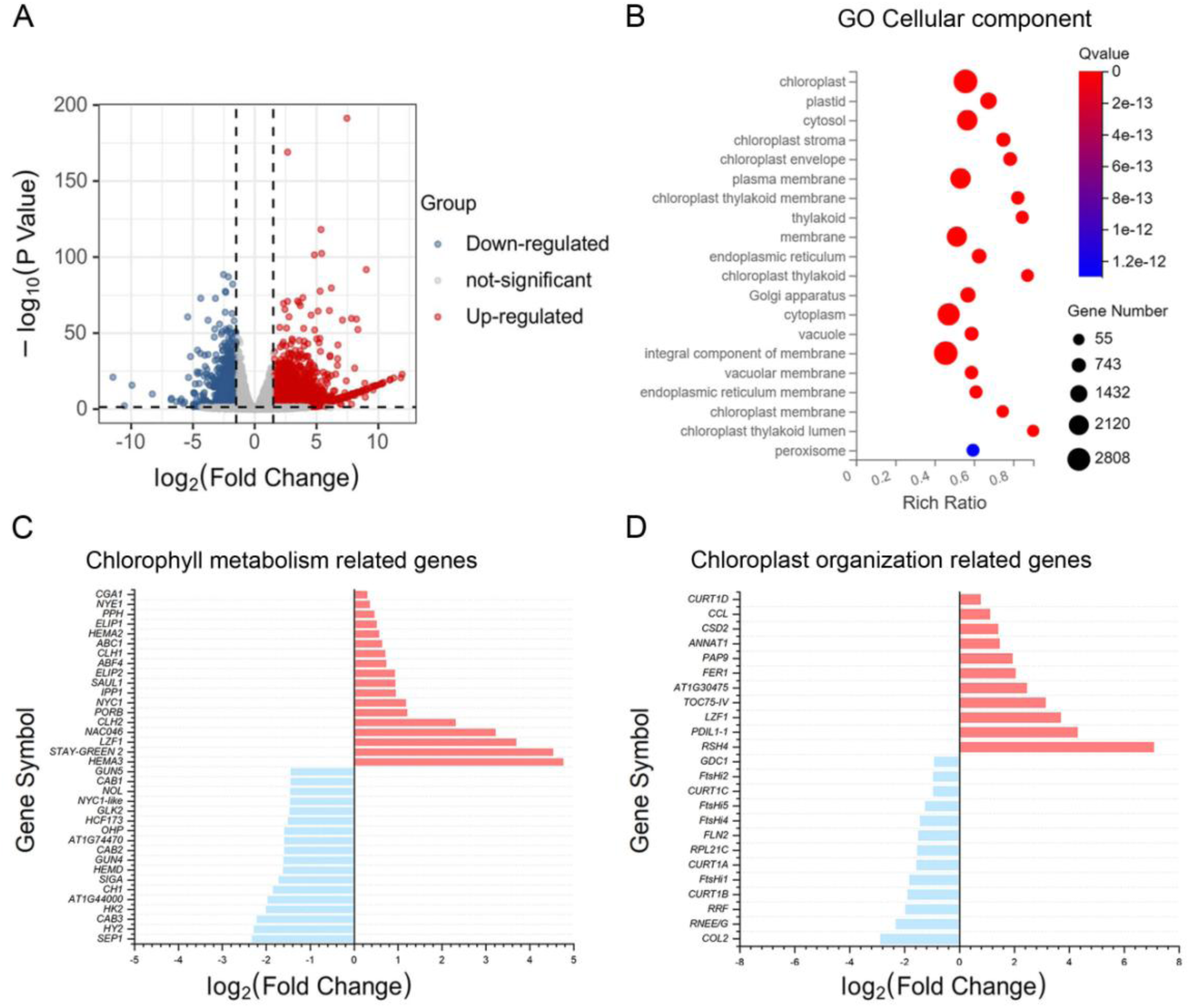
Transcriptomic analysis reveals impaired expression of chloroplast-related genes in the *hda6* mutant. (A) Volcano plot showing differential gene expression between *axe1-4* and the WT plants. Up-regulated and down-regulated genes are indicated in red and blue, respectively, while non-significant genes are shown in gray. (B) Gene Ontology (GO) enrichment analysis of differentially expressed genes in the cellular component category. Significantly enriched terms are mainly associated with chloroplast-related compartments, including the chloroplast envelope, thylakoid, and stroma. The color gradient represents the *q*-value, and bubble size corresponds to the number of genes within each category. (C) Differential expression of genes involved in chlorophyll metabolism. Several key genes related to chlorophyll biosynthesis are down-regulated in *axe1-4*. (D) Differential expression of genes associated with chloroplast organization and biogenesis. Genes encoding structural and functional components of the chloroplast are predominantly suppressed in *axe1-4* compared with WT.

Such repression mirrors the genome-wide reduction in photosynthetic and plastidic transcripts reported in *hda6* and *hda9* mutants, where altered chromatin acetylation leads to transcriptional silencing of organelle-related gene clusters. HDA6 has been shown to physically associate with MET1 and LDL1/2 to coordinate histone deacetylation and DNA methylation, thereby maintaining stable gene expression patterns (Liu *et al*., 2012). It also functions with FLD to repress FLC, demonstrating that HDA6 participates in multiprotein complexes that integrate histone modification and transcriptional regulation (Wu *et al*., 2008).

Furthermore, HDA6-dependent deacetylation has been implicated in mRNA 3’-end processing and polyadenylation site selection, suggesting that its loss could simultaneously affect transcript stability and splicing (Lin *et al*., 2020). In the context of chloroplast-related genes, such post-transcriptional mis-processing could amplify transcriptional repression, leading to global down-regulation of the chloroplast proteome. Collectively, these results indicate that HDA6 may act at multiple regulatory levels including chromatin remodeling, transcriptional activation and RNA maturation, to coordinate nuclear control of chloroplast development.

### HDA6 promotes the assembly and stability of photosynthetic complexes

The integrity of thylakoid protein assemblies was next evaluated using Blue Native PAGE (BN-PAGE). As shown in Figure 3A, thylakoid membranes from *axe1-4* accumulated reduced levels of PSI, PSII, cytochrome b₆f, and LHCII supercomplexes relative to the wild type. Subsequent two-dimensional SDS-PAGE (Figure 3B) resolved subunit composition, revealing a marked decrease in PSI (PsaA–E), PSII (PsbA–D), ATP synthase (AtpA–C), and LHC (Lhca1–3, Lhcb1–5) proteins. These reductions suggest defective assembly or enhanced turnover of photosynthetic pigment-protein complexes.

**Figure 3.**
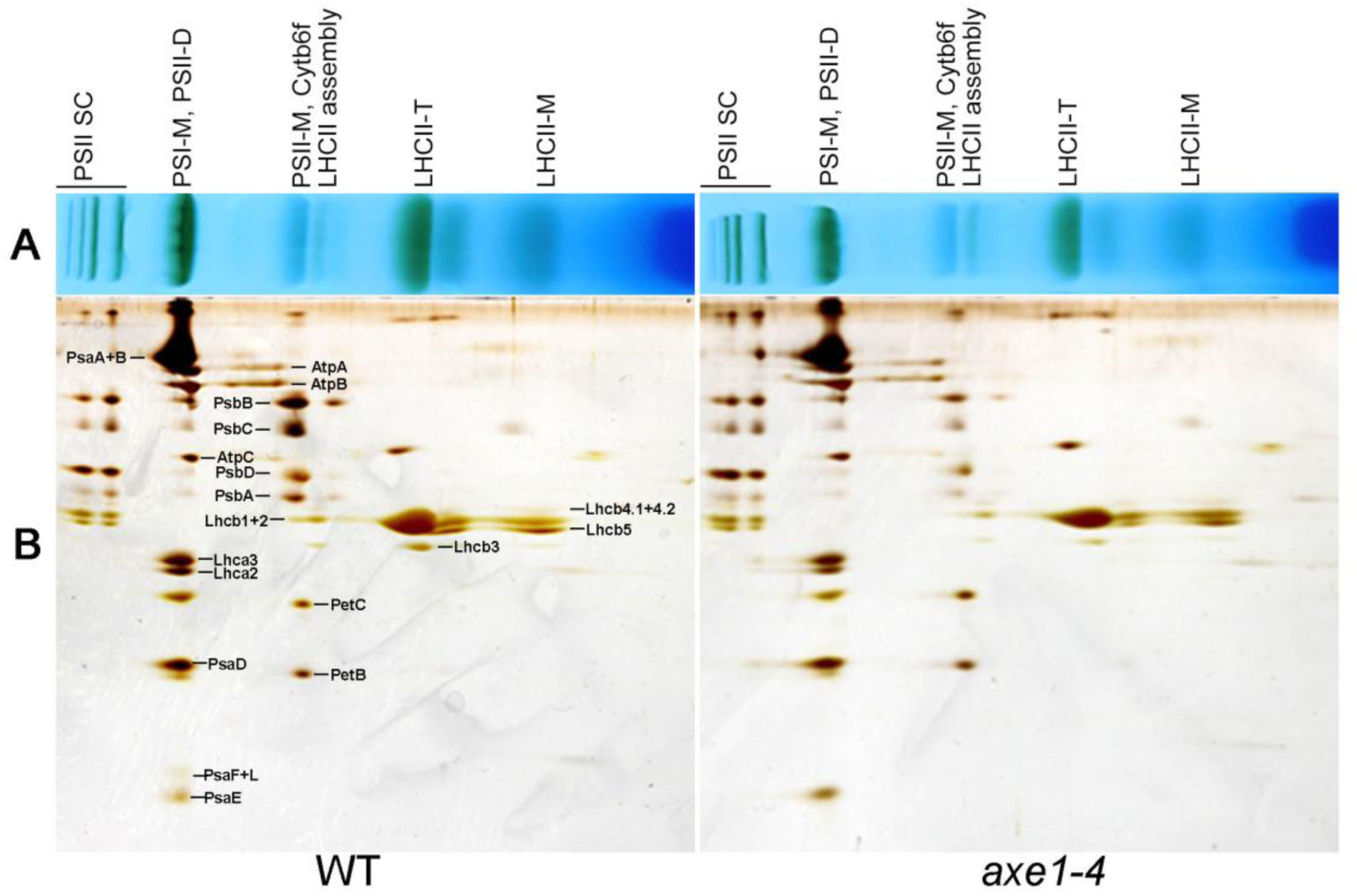
Loss of HDA6 reduces the assembly of photosynthetic protein complexes. (A) Blue Native-PAGE (BN-PAGE) analysis of thylakoid membrane protein complexes isolated from WT and *axe1-4* mutants. The major photosynthetic supercomplexes, including PSII supercomplex (PSII SC), PSI monomer (PSI-M), PSII dimer (PSII-D), PSII monomer (PSII-M), cytochrome b₆f (Cytb₆f), and light-harvesting complex II (LHCII) trimer and monomer (LHCII-T, LHCII-M) are indicated. (B) Two-dimensional BN/SDS–PAGE separation and coomassie staining of thylakoid membrane proteins from WT and *axe1-4*. Subunits of PSI (PsaA-E, PsaF+L), PSII (PsbA-D), ATP synthase (AtpA-C), Cytb₆f (PetB, PetC), and light-harvesting complexes (Lhca1-3, Lhcb1-5) are annotated. The *axe1-4* mutant shows a marked reduction in the accumulation of PSI, PSII, and LHCII components compared with WT, indicating defective assembly or stability of photosynthetic protein complexes.

Comparable phenotypes are seen in *curt1* and *alb3* mutants, in which altered thylakoid curvature or protein insertion reduces photosystem supercomplex stability (Armbruster *et al*., 2013; Ackermann *et al*., 2021). Given that many subunits of PSI, PSII, and ATP synthase undergo reversible lysine acetylation, HDA6 may indirectly influence their folding or membrane integration by modulating acetylation of nuclear-encoded chloroplast proteins. Proteomic studies have revealed that chloroplast enzymes such as Rubisco activase and ATP synthase subunits are dynamically acetylated and their deacetylation status correlates with catalytic efficiency and complex stability (Finkemeier *et al*., 2011). Hence, *HDA6* mutation may lead to hyperacetylation of these proteins, disrupting photosystem stoichiometry and impairing energy conversion.

### HDA6 is required for maintaining thylakoid architecture and grana stacking

Transmission electron microscopy provided ultrastructural insight into the observed physiological impairments. Figure 4A illustrates the pronounced differences between the WT and *axe1-4* chloroplasts. The chloroplasts in the WT possessed tightly stacked grana interconnected by stroma lamellae, whereas *axe1-4* displayed swollen thylakoid membranes, decreased grana height and irregular lamellar spacing. Quantitative measurements confirmed significant reductions in both the number of thylakoids per granum and average membrane thickness. Complementary cryo-electron tomography (cryo-ET) revealed three-dimensional disorganization, with the mutant lacking the regular vertical alignment typical of mature grana stacks (Figure 4B).

**Figure 4.**
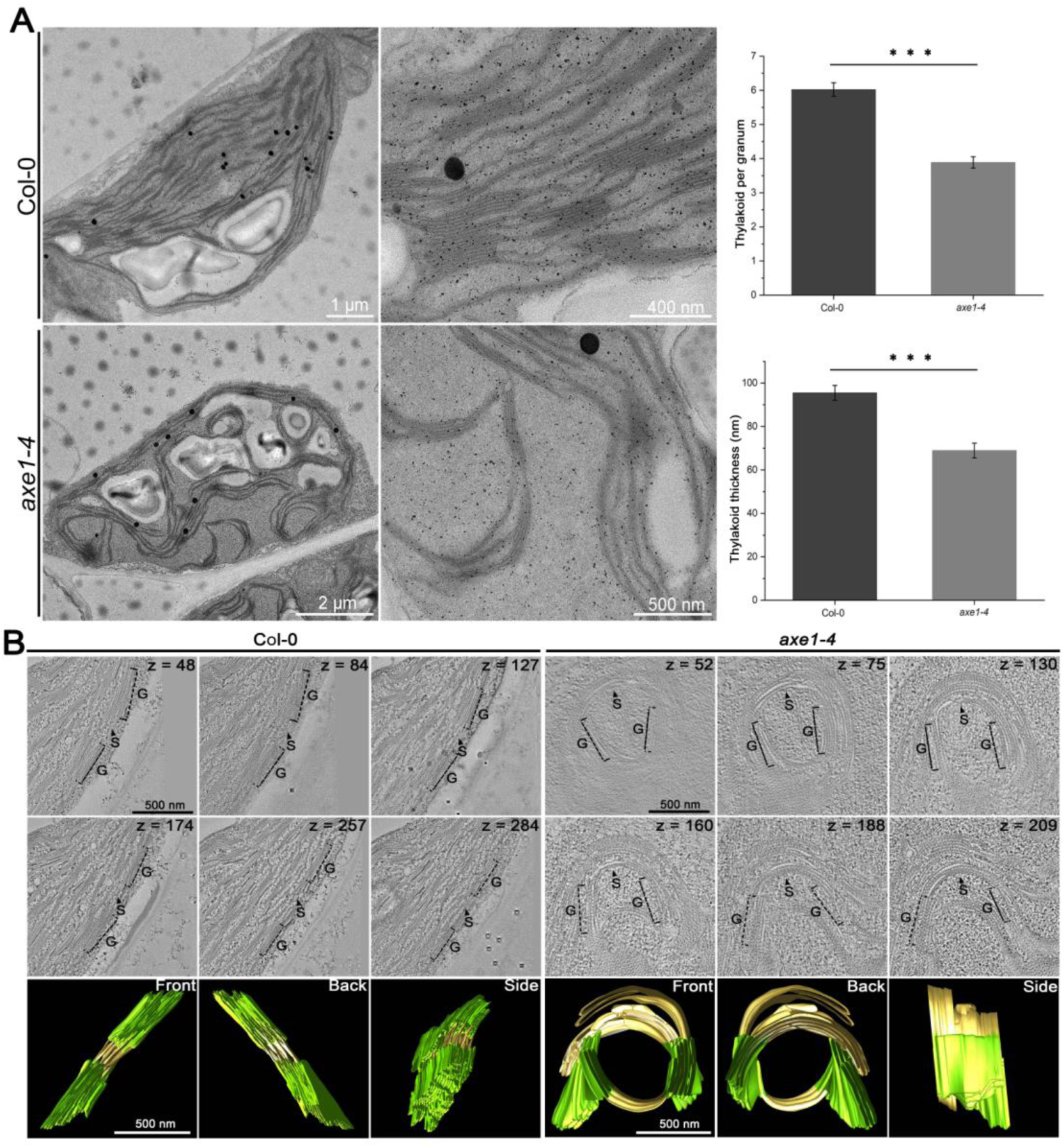
HDA6 is essential for maintaining proper thylakoid architecture and grana stacking in chloroplasts. (A) Transmission electron microscopy (TEM) images showing chloroplast ultrastructure in mesophyll cells of WT and *axe1-4* mutants. The *axe1-4* mutant exhibits distorted thylakoid membranes and loosely stacked grana compared with the well-organized grana stacks observed in WT. Quantitative analyses (right) show significant reductions in the number of thylakoids per granum and in thylakoid membrane thickness in *axe1-4* relative to WT (mean ± SD; \*\**p* < 0.001, Student’s *t*-test). (B) Representative images of cryo-electron tomography (Cryo-ET) ultrastructural study of chloroplasts from WT and *axe1-4* mesophyll cells. Sequential z-slice images depict the internal organization of grana (G) and stroma lamellae (S). Three-dimensional renderings (bottom) reveal the compact and orderly grana stacks in WT compared with the disorganized and irregular thylakoid arrangements in *axe1-4*. Scale bars are indicated in each panel.

These defects resemble those of *vipp1* and *mgd1* mutants, in which disruption of membrane biogenesis or galactolipid synthesis compromises thylakoid formation (Nordhues *et al*., 2012; Rocha *et al*., 2016). Recent tomographic analyses have shown that proper thylakoid stacking depends on curvature-inducing protein1 (CURT1) and precise lateral heterogeneity between PSII-rich grana and PSI-enriched stroma lamellae (Dann *et al*., 2025). The resemblance of *axe1-4* to these structural mutants suggests that HDA6 indirectly governs thylakoid morphogenesis through transcriptional regulation of key architectural genes or through deacetylation of chloroplast proteins that stabilize membrane curvature. By maintaining thylakoid ultrastructure, HDA6 ensures optimal photosynthetic efficiency and energy distribution within the chloroplast.

### HDA6 partially localizes to chloroplasts and functions as a non-histone deacetylase

Subcellular localization of HDA6 was examined using *pUBQ::HDA6-GFP* transgenic Arabidopsis. Confocal microscopy revealed predominant GFP fluorescence in the nucleus and cytoplasm (Figure 5), with partial overlap with chlorophyll autofluorescence inside chloroplasts (Figure 5). This dual localization implies that HDA6 operates in both nuclear and organellar compartments. Similar nucleocytoplasmic partitioning has been observed for Class II HDACs such as AtHDA14 and OsHDA710, which are targeted to mitochondria and chloroplasts to deacetylate metabolic enzymes involved in the tricarboxylic acid cycle and photosynthesis (Liu *et al*., 2022).

**Figure 5.**
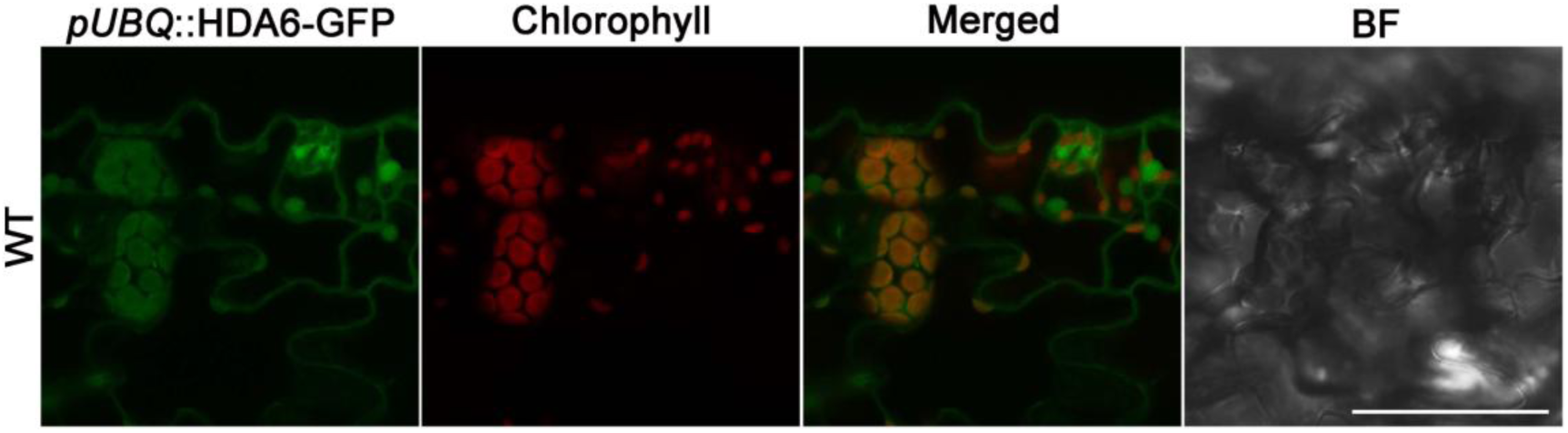
HDA6 localizes in Arabidopsis chloroplasts. Confocal laser scanning microscopy of *pUBQ::HDA6-GFP* transgenic Arabidopsis plants. GFP fluorescence (green) represents the localization of HDA6-GFP, while chlorophyll autofluorescence (red) indicates the position of chloroplasts. Merged images show that HDA6-GFP typically localizes in the nucleus and cytoplasm, as well as overlap with chlorophyll signals, suggesting that HDA6 also localizes in leaf chloroplasts. The bright-field (BF) image provides a structural view of the epidermal cells. Scale bar = 20 μm.

The chloroplast-localized fraction of HDA6 may act directly on non-histone substrates, regulating redox homeostasis and photosynthetic protein turnover. Previous biochemical analyses of rice HDACs showed that several isoforms physically associate with stromal proteins and modulate their acetylation state under light stress conditions (Li *et al*., 2021). Hence, HDA6 might contributes to chloroplast proteostasis, coupling nuclear epigenetic regulation with local post-translational control inside the organelle.

Our findings establish HDA6 as a multifunctional regulator that integrates epigenetic, transcriptional, and metabolic control of chloroplast development. Beyond its canonical function in flowering and senescence (Wu *et al*., 2008). HDA6 coordinates nuclear gene expression with chloroplast morphogenesis and energy metabolism. The *axe1-4* mutant phenotype, encompassing chlorophyll loss, reduced photosystem assembly, and defective thylakoid structure, reflects disruption of both nuclear transcriptional networks and organellar protein acetylation. We propose a hypothetical model in which HDA6 acts in two interconnected layers: i) in the nucleus, HDA6 deacetylates histones at promoters of chloroplast-associated genes, promoting their transcriptional activation might in coordination with MET1, LDL1/2, and FLD complexes; and ii) within chloroplasts, HDA6 deacetylates non-histone proteins to preserve photosynthetic protein stability and membrane organization. Through these dual mechanisms, HDA6 serves as a molecular bridge linking chromatin dynamics with chloroplast proteome homeostasis.

Collectively, our results demonstrate that HDA6 is an essential regulator of chloroplast biogenesis and photosynthetic function in Arabidopsis. Loss of HDA6 disrupts chlorophyll accumulation, down-regulates plastid gene expression, impairs photosystem complex assembly, and destabilizes thylakoid architecture. These findings expand the known functional repertoire of plant histone deacetylases, revealing an evolutionarily conserved strategy by which chromatin modifiers orchestrate organelle development and energy metabolism. HDA6 therefore represents a key node in the epigenetic network that synchronizes nuclear transcription with chloroplast differentiation, ensuring efficient photosynthetic performance and plant vitality.

## Materials and Methods

### Plant materials and growth conditions

Arabidopsis thaliana ecotype Columbia-0 (Col-0) was used as the WT control in all experiments. The *axe1-4* mutant, carrying a loss-of-function allele of HDA6 was previously described (Earley *et al*., 2006). Arabidopsis seeds were surface-sterilized with 75% (v/v) ethanol for 10 min and rinsed with sterile water for three times, and sown on ½ Murashige and Skoog (MS) salts supplemented with 1% sucrose, pH 5.7, and 0.8% (w/v) agar in Petri dishes. Thereafter, the plates with sterilized seeds were stratified at 4 °C for 2 d and transferred in a plant growth chamber for 7 d prior to being grown in soil in a green house at 22 °C, 40-50% relative humidity and 80 µmol m^−2^ • s^−1^ light intensity under long-day (16 h light and 8 h dark) conditions.

### Measurement of chlorophyll content

Chlorophyll a and b contents were quantified as previously described (Porra *et al*., 1989). 50 mg fresh 30-day-old leaves were homogenized in 1 mL of 80% (v/v) acetone, followed by centrifugation at 12,000 × g for 10 min at 4 °C. Absorbance of the supernatant was measured at 663 nm and 646 nm using a UV–visible spectrophotometer (Bio-Rad), and chlorophyll concentrations were calculated according to standard equations. Three biological replicates were analyzed per genotype.

### Confocal microscopy and chloroplast imaging

Leaf epidermal and mesophyll cells from fully expanded leaves were examined using Leica TCS SP8 system with the following parameters: 63 × oil objective, 2 × zoom, 750 gain, 0 background, 0.168 µm pixel size and photomultiplier tubes detector. Chlorophyll autofluorescence was excited at 488 nm and detected at 650–750 nm. For subcellular localization, GFP signals in *pUBQ::HDA6-GFP* transgenic plants were visualized under identical settings and compared with chlorophyll autofluorescence. Chloroplast size and morphology were quantified from confocal images using ImageJ (NIH), with at least 50 chloroplasts analyzed from each of three independent plants.

### Transmission electron microscopy and cryo-electron tomography

Ultrastructural analysis of chloroplasts was performed following standard fixation and embedding protocols. Leaf segments (1 mm²) were fixed in 2.5% (v/v) glutaraldehyde and 1% (w/v) paraformaldehyde in 0.1 M phosphate buffer (pH 7.2) for 2 h, post-fixed in 1% (w/v) osmium tetroxide, dehydrated through a graded ethanol series, and embedded in Spurr’s resin. Ultrathin sections (70 nm) were stained with uranyl acetate and lead citrate, and examined using a JEOL JEM-1400 transmission electron microscope at 80 kV. Grana number, thylakoid width, and membrane stacking parameters were quantified using ImageJ. For cryo-electron tomography, leaf samples were plunge-frozen in liquid ethane and examined using a Thermo Fisher Talos Arctica cryo-TEM equipped with a Falcon III camera. Tomograms were reconstructed using IMOD software, and three-dimensional renderings of thylakoid membranes were generated to assess grana organization and membrane continuity.

### Transcriptomics analysis

For the transcriptomics analysis, 30-day-old leaves of WT and *axe1-4* were collected and sent to BGI (https://www.bgi.com/) for high-throughput sequencing using the Illumina platform. For each sample, 2 µg of total RNA was used as input material for RNA library preparation. The sequencing libraries were generated using the NEBNext Ultra RNA Library Prep Kit for Illumina (Catalog No. E7530L, NEB), following the manufacturer’s recommended protocol. To facilitate sample identification during downstream analyses, unique index codes were incorporated into each library.

### Blue-native PAGE and two-dimensional protein complex analysis

Thylakoid membranes were isolated from 4-week-old leaves as previously described (Ouyang *et al*., 2020). Chlorophyll concentration was adjusted to 1 mg mL⁻¹, and membrane protein complexes were solubilized with 1% (w/v) n-dodecyl β-D-maltoside. Native protein complexes were separated by 4–12% Blue Native PAGE and visualized by Coomassie staining. For second-dimension separation, individual lanes were excised, incubated in SDS sample buffer, and resolved on 12% SDS-PAGE gels. Gels were stained with silver staining.

### Statistical analysis

All experiments were performed with at least three independent biological replicates. Statistical significance was asses sed using Student’s *t*-test or one-way ANOVA with Tukey’s post hoc test, as appropriate. Data are presented as means ± standard deviation (SD). Differences were considered significant at *P* < 0.05.

## Acknowledgements

The author sincerely apologizes for any shortcomings and limitations that remain, as this manuscript was prepared within a constrained timeframe following the unauthorized use and theft of certain results by an individual not involved in this study. This work is supported by grants from the National Natural Science Foundation of China (91954110) and the Natural Science Foundation of Guangdong Province (2021A1515012066) to H.W.

## Author contributions and competing interests

H.W. conceived and designed the study, performed the experiments, analyzed the data, prepared the figures, wrote the manuscript and obtained funding support. The author declares no competing interests.

